# Nanopore- and AI-empowered microbial viability inference

**DOI:** 10.1101/2024.06.10.598221

**Authors:** Harika Ürel, Sabrina Benassou, Hanna Marti, Tim Reska, Ela Sauerborn, Yuri Pinheiro Alves De Souza, Albert Perlas, Enrique Rayo, Michael Biggel, Stefan Kesselheim, Nicole Borel, Edward J. Martin, Constanza B. Venegas, Michael Schloter, Kathrin Schröder, Jana Mittelstrass, Simone Prospero, James M. Ferguson, Lara Urban

## Abstract

The ability to differentiate between viable and dead microorganisms in metagenomic data is crucial for various microbial inferences, ranging from assessing ecosystem functions of environmental microbiomes to inferring the virulence of potential pathogens from metagenomic analysis. While established viability-resolved genomic approaches are labor-intensive as well as biased and lacking in sensitivity, we here introduce a new fully computational framework that leverages nanopore sequencing technology to assess microbial viability directly from freely available nanopore signal data. Our approach utilizes deep neural networks to learn features from such raw nanopore signal data that can distinguish DNA from viable and dead microorganisms in a controlled experimental setting of UV-induced *Escherichia* cell death. The application of explainable AI tools then allows us to pinpoint the signal patterns in the nanopore raw data that allow the model to make viability predictions at high accuracy. Using the model predictions as well as explainable AI, we show that our framework can be leveraged in a real-world application to estimate the viability of obligate intracellular *Chlamydia*, where traditional culture-based methods suffer from inherently high false negative rates. This application shows that our viability model captures predictive patterns in the nanopore signal that can be utilized to predict viability across taxonomic boundaries. We finally show the limits of our model’s generalizability through antibiotic exposure of a simple mock microbial community, where a new model specific to the killing method had to be trained to obtain accurate viability predictions. While the potential of our computational framework’s generalizability and applicability to metagenomic studies needs to be assessed in more detail, we here demonstrate for the first time the analysis of freely available nanopore signal data to infer the viability of microorganisms, with many potential applications in environmental, veterinary, and clinical settings.

**Author summary:** Metagenomics investigates the entirety of DNA isolated from an environment or a sample to holistically understand microbial diversity in terms of known and newly discovered microorganisms and their ecosystem functions. Unlike traditional culturing of microorganisms, genomic approaches are not able to differentiate between viable and dead microorganisms since DNA might persist under different environmental circumstances. The viability of microorganisms is, however, of importance when making inferences about a microorganism’s metabolic potential, a pathogen’s virulence, or an entire microbiome’s impact on its environment. As existing viability-resolved genomic approaches are labor-intensive, expensive, and lack sensitivity, we here investigate our hypothesis if freely available nanopore sequencing signal dat that captures DNA molecule information beyond the DNA sequence might be leveraged to infer such viability. This hypothesis assumes that DNA from dead microorganisms accumulates certain damage signatures that reflect microbial viability and can be read from nanopore signal data using fully computational frameworks. We here show first evidence that such a computational framework might be feasible by training a deep model on controlled experimental data to predict viability at high accuracy, exploring what the model has learned, and using it in a real-world application by application to a bacterial species of veterinary relevance. We finally show that a specific model has to be trained to accurately predict viability after antibiotic exposure of a mock microbial community. While the generalizability of our computational framework therefore needs to be assessed in much more detail, we here demonstrate that freely available data might be usable for relevant viability inferences in environmental, veterinary, and clinical settings.

## Introduction

While microbial cultivation remains a foundational technique in microbiology to assess the taxonomic composition of microbial communities and to understand their physiology and ecosystem functions [1], only a small fraction of microbial diversity has been isolated in pure culture [2]. This limitation has led to undiscovered functions and biased representations of the phylogenetic diversity of microbial communities in nearly all of Earth’s environments [2]. While medically relevant microorganisms of the human microbiome often constitute an exemption since they have been disproportionately well studied through microbial cultures [3], the clinical application of microbial cultivation for pathogen profiling is further limited by its time-consuming and labor-intensive nature [4].

The first studies of the so-called “microbial dark matter” have been enabled by advances in culture-independent molecular methodology [5], and have been based on amplifications of conserved marker regions such as ribosomal RNA genes [6]. Such targeted metabarcoding approaches, however, suffer from several limitations: They can often not provide strain- or even species-level taxonomic resolution, are highly dependent on genomic database completeness, do not allow for any functional inferences or virulence annotations, and often introduce amplification bias due to differential amplification efficiency and primer mismatches, which can significantly distort the representation of microbial community compositions [7].

Metagenomics, on the other hand, is a shotgun sequencing-based molecular methodology that can assess the entirety of DNA isolated from an environment or a sample and *de novo* assemblies of potentially complete microbial genomes of all present microorganisms; such genome-based approaches provide a variety of phylogenetically informative sequences for taxonomic classification, information about the metabolic and virulence potential of microorganisms, and the potential to identify completely novel genes [8, 9].

Especially long-read metagenomic approaches have shown great promise in achieving highly contiguous *de novo* assemblies through the recovery of high-quality metagenome-assembled genomes (MAGs) from complex environments; specifically, the latest advances in nanopore sequencing technologies have resulted in high sequencing accuracies of very long sequencing reads of up to millions of bases, which allowed for the generation of hundreds of MAGs from metagenomic data, including the generation of closed circularized genomes [10, 11]. Nanopore sequencing technology is based on the interpretation of the disruption of an ionic current due to a motor protein guiding individual nucleotide strands through nanopores embedded in an electrically resistant polymer membrane at a consistent translocation speed [12]. This raw nanopore signal, or “squiggle” data, can then be translated into nucleotide sequence using bespoke neural network-based basecalling algorithms [13], which—when efficiently embedded on powerful GPUs—can generate genomic data in real-time. The portable character and straightforward implementation of nanopore sequencing at low upfront investment costs further make this technology accessible for fast microbial and pathogen assessments at the point of interest all around the world, including in low- and middle-income countries [14].

In contrast to cultivation-based approaches, molecular methods suffer from their inherent deficiency of not being able to differentiate between viable and dead microorganisms [2, 15]. While cultivation-based approaches only detect viable microorganisms, DNA might remain intact and therefore accessible by molecular methods despite the respective microorganisms being dead [15, 16]. This would be relevant in the context of clinical infection prevention and control and pathogen monitoring, where certain disinfection methods or the use of systemic antibiotics often kill the bacteria before the DNA is destroyed [15, 17], but also for understanding the ecosystem functions of thus far understudied microbiomes [2]: For example, the air microbiome has been shown to be remarkable diverse and variable when assessed through nanopore metagenomics [18], but given the low biomass of this environment it is expected that many microorganisms might be dead and stem from adjacent environments such as soil or water. The persistence of the DNA of dead microorganisms in the environment might hereby depend on many factors, including external conditions such as temperature, pH, and microbial activity, and internal, taxon-specific parameters such as microbial cell wall composition. Viability-resolved metagenomics would, however, be crucial for the interpretation of metagenomic data, ranging from outbreak source detection [19], food safety [20], and public health investigations [21], to ecosystem function inferences [22].

To assess microbial viability from genomic data, several approaches have been developed: Culture-dependent viability methods combine the advantages of cultivation-based and molecular approaches by growing certain microorganisms of interest on selective media; this approach, however, remains time-consuming and labor-intensive and suffers from the same selectivity of growth media and culturable microorganisms as purely cultivation-based approaches [23], especially for fastidious or obligate intracellular microorganisms [24]. Microbial viability has further been described by metabolic activity, where microbial cells are incubated with specific substrates leading to ATP production, tetrazolium salt reduction, or radiolabeled substrate incorporation [25]. Further, ribosomal RNA may be assessed as a read-out of microbial activity [26]. To what extent such metabolic activity can be used as a proxy for microbial viability, however, remains to be explored [25]. While messenger RNA has been used as a viable/dead marker due to its intrinsic instability outside of the microbial cell [27, 28], the metatranscriptome still has to be stable enough in the environment to be detectable at all, potentially leading to many false-negative detections; if only one gene is targeted, the analyzed gene further has to be expressed shortly before cell death. Additional potential problems stem from the relatively challenging extraction protocols due to the RNA’s instability and from the evolutionary conservation of gene sequences, which can hamper taxonomic resolution [15].

Finally, an aspect that can be used for viability-resolved metagenomics is the physical difference between viable and dead cells: Viability PCR (vPCR) uses DNA-intercalating dyes such as ethidium monoazide (EMA) or propidium monoazide (PMA) to differentiate between viable and dead cells. These dyes penetrate only dead cells with compromised membranes and bind to their DNA via covalent bonds upon photoactivation, preventing it from being amplified during subsequent PCR [15, 17]; this approach has been applied to a diverse array of Gram-negative and -positive bacteria, and to assess the effectiveness of disinfection and heat treatment [25]. It, however, relies on the assumption that membrane integrity is a reliable indicator of viability, which can lead to overestimation of viability if cells lose viability without immediate membrane compromise [29] and can be biased by the dye’s variable permeability across different microbial cell wall structures [30, 31]. The dependence of the approach on photoactivation further means that turbid material might hamper the efficiency of the dye [32].

All these established viability-resolved metagenomic approaches are labor-intensive, require additional reagents and sample processing, and are often biased and lack sensitivity. We here hypothesized that the raw, freely available nanopore signal from metagenomic datasets might be leveraged to infer microbial viability, assuming that the native DNA from dead microorganisms accumulates detectable squiggle signatures due to, e.g., external damage through UV, heat, or drought exposure, the lack of DNA repair mechanisms, or enzymatic degradation activity [33, 34, 35]. Such an analysis framework could be fully computational and utilize squiggle data that is automatically obtained with nanopore sequencing. While raw nanopore data is known to contain information about epigenetic modifications [36, 37, 38] and oxidative stress at specific human telomere sites [39], the applicability to assess microbial viability has not yet been tested.

In this study, we produced experimental nanopore sequencing data from viable and UV-killed *Escherichia coli* cultures to optimize deep neural networks to predict viability just from the nanopore squiggle signal. We then applied explainable AI (XAI) tools, which allow us to identify the specific nanopore signal patterns in the input data that allow the model to deliver high-accuracy predictions as an output. We show that our computational framework can be leveraged in a real-world application to estimate the viability of obligate intracellular *Chlamydia suis*, pointing towards the applicability of our model across taxonomic boundaries, including to species with highly complex life cycles. We finally explore the limits of our model’s generalizability through antibiotic exposure of a simple mock microbial community, where we had to train a new killing method-specific model to obtain accurate viability predictions. While the extent of our computational framework’s generalizability needs to be assessed in more detail, we here demonstrate for the first time the potential of analyzing freely available nanopore signal data to infer the viability of microorganisms, with many applications in environmental, veterinary, and clinical settings.

## Results & Discussion

### Viability model training and inference

We generated controlled training data by nanopore sequencing native DNA of viable and dead *E. coli* (Materials and Methods). We killed *E. coli* cultures using different stressors to then isolate the extracellular DNA and expose it to natural degradation. We only obtained enough DNA for subsequent shotgun sequencing from the viable culture and from the culture killed through rapid UV exposure (viable: 212 ng/μL; UV: 5.46 ng/μL; heat shock: 0.03 ng/μL; bead beating: 0.67 ng/μL; Materials and Methods). We repeated this experiment and confirmed that rapid heat shock, as well as bead beating exposure, again resulted in very low DNA concentrations, suggesting quick and complete DNA degradation. We hypothesize that UV exposure is the only stressor in our study that simultaneously destroys bacterial cell walls and inactivates DNA-degrading enzymes. In contrast, heat shock at 120°C and bead beating might not uniformly degrade all enzymatic activity [40, 41], potentially allowing residual DNA-degrading enzymes to persist and contribute to the degradation of genomic material during subsequent natural exposure. We therefore created nanopore shotgun sequencing of the viable and the UV-exposed culture, which resulted in 2.92 Gbases (Gb; median read length of 2,476 b) and 2.69 Gb (median read length of 1,606 b) of sequencing output, respectively (Materials and Methods).

We then tested the implementation of different neural network architectures to predict the binary viability state from the raw nanopore data (0=viable; 1=dead after UV exposure; Materials and Methods). We processed the *E. coli* nanopore signal, or “squiggle”, data, cut it into altogether 3,181,600 signal chunks of 10k signals, and separated the chunks into balanced training (60%), validation (20%), and test (20%) set along each original sequencing read to avoid that signal chunks from the same read would end up in the same dataset (Materials and Methods). These signal chunks were then treated as 1D time series signal data of consistent length. We trained the different model architectures using different learning rates (LRs) up to 1,000 epochs, assessing the models’ performance based on training and validation loss after each epoch (Materials and Methods; **Table S1**; **Fig S1**). The loss plot of our best-performing model, a residual neural network with convolutional input layers (configuration ResNet1; LR=1e-4; **Table S1**; **Fig S1**; Materials and Methods) shows minimal overfitting when the minimum validation loss is reached at epoch 667 (**Fig 1A**). The other residual neural network architectures (ResNet2, ResNet3), on the other hand, resulted in overfitting to the training data at any LR, and the transformer architecture did not reach the minimum validation loss of ResNet1 (**Fig S1**). We next only focused on ResNet1 and optimized its probability threshold using the validation set; in order to obtain a high accuracy, we maintained the probability threshold at the default value of 0.5 (**Fig 1B**), which resulted in a good final performance on the test data with an accuracy of 0.83 and a F1 score of 0.81 (**Fig 1B**, *inlet*) as well as Area Under the Curve (AUC) values of 0.90 (Area Under the Receiver Operating Characteristic curve; AUROC) and 0.92 (Area Under the Precision-Recall curve; AUPR; **Fig 1C**).

**Fig 1.**
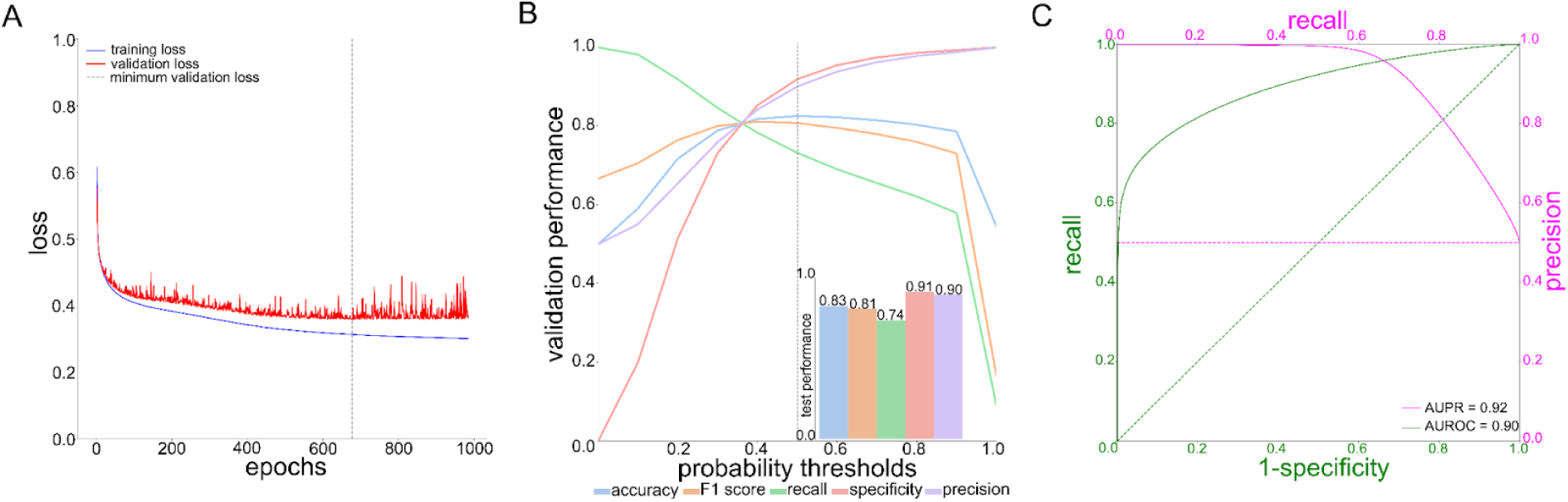
**Training and viability inference on UV-killed *E. coli* of the Residual Neural Network 1 (ResNet1)** (A) Model loss for training and validation datasets across 1,000 epochs; the minimum validation loss of ResNet1 was reached at epoch 677. (B) Prediction probability threshold optimization on the validation dataset resulted in a probability threshold of 0.5 for obtaining maximum accuracy. *Inlet:* Performance of ResNet1 on the test dataset (Materials and Methods). (C) Test dataset performance of ResNet1 in terms of Precision-Recall (PR; *magenta)* and Receiver Operating Characteristic (ROC; *green*) curves and their respective Areas Under the Curve (AUPR, AUROC).

We also trained the same residual neural network architecture ResNet1 on the basecalled nanopore data of viable and dead *E. coli* at a standardized chunk size of 800 b, which roughly corresponds to the signal chunk size of 10k signals (Materials and Methods). Independent of whether we only basecalled the canonical bases or used a N6-methyladenine (6mA) modification-aware basecalling model (Materials and Methods), the model could not be trained to distinguish viable from dead data just from basecalled DNA sequence data (**Table S1**). This shows that our model captures patterns in the squiggle data that go beyond the encoding of nucleotides and their known epigenetic modifications. While this was expected since we used the same *E. coli* culture with the same reference genome to create the viable and dead datasets, we can rule out that our squiggle-based model captured any random differences in DNA sequence context between the two datasets that might have occurred by chance.

We additionally obtained the performance of ResNet1 for different signal chunk sizes (**Fig S2**; **Table S1**), and we found that viability prediction performance was possible from a minimum chunk size of approximately 5k, but that performance further improved with increasing chunk size. This shows that larger signal chunks contain more information that can be used by our model to make more accurate per-chunk predictions despite a consequently reduced size of the training dataset. We here stick to our original model with a chunk size of 10k signals, which resulted in relatively good performance (**Fig 1**; **Table S1**).

We finally trained two logistic regression models using only the read length and translocation speed per sequencing read as input feature, respectively, to compare the performance of our squiggle-based deep neural network with simple baseline models. Both models substantially underperformed in comparison to our model (**Table S1**). The read length model performed similarly to a random classifier (accuracy of 0.5; **Table S1**), showing that the read length distribution difference between the viable and dead sequencing datasets can not be leveraged to infer viability in this dataset. We also argue that while a difference in read length distribution between viable and dead microbes might be expected in other settings, for example after stronger or longer degradation exposure, such a difference would have to be substantial to allow for accurate viability classifications of each individual sequencing read. In mixed microbial communities, the read length distribution would further be confounded by the microbial composition and the respective genome sizes. The translocation speed model reached a slightly higher accuracy of 0.59 (**Table S1**), which might be explained by UV-induced twists or kinks in the DNA backbone having a slight impact on the translocation of the sequencing read.

### Explainable AI application

We implemented Class Activation Maps (CAM) as an XAI method [42] to identify the most important regions in the nanopore signal data that inform the model’s viability classifications (Materials and Methods; **Fig 2A**). We found that “dead” signal chunks exhibited discrete regions of increased CAM values (“CAM regions” defined at CAM values>0.8; **Fig 2B** for several true positive classifications of the test dataset). To confirm the importance of these CAM regions for the model’s final predictions, we applied consecutive masking of the regions with the highest CAM values within each nanopore signal chunk (**Fig S3** for several examples); we observed that the prediction probability for being classified as “dead” decreased with increased masking of CAM-relevant regions, either by consecutively masking regions using a consistent mask size or by increasing the mask size (from 100 to 2k signals; **Fig 2C**). This shows that the CAM application reliably pinpoints patterns in the nanopore signal that are predictive for our viability model.

**Fig 2.**
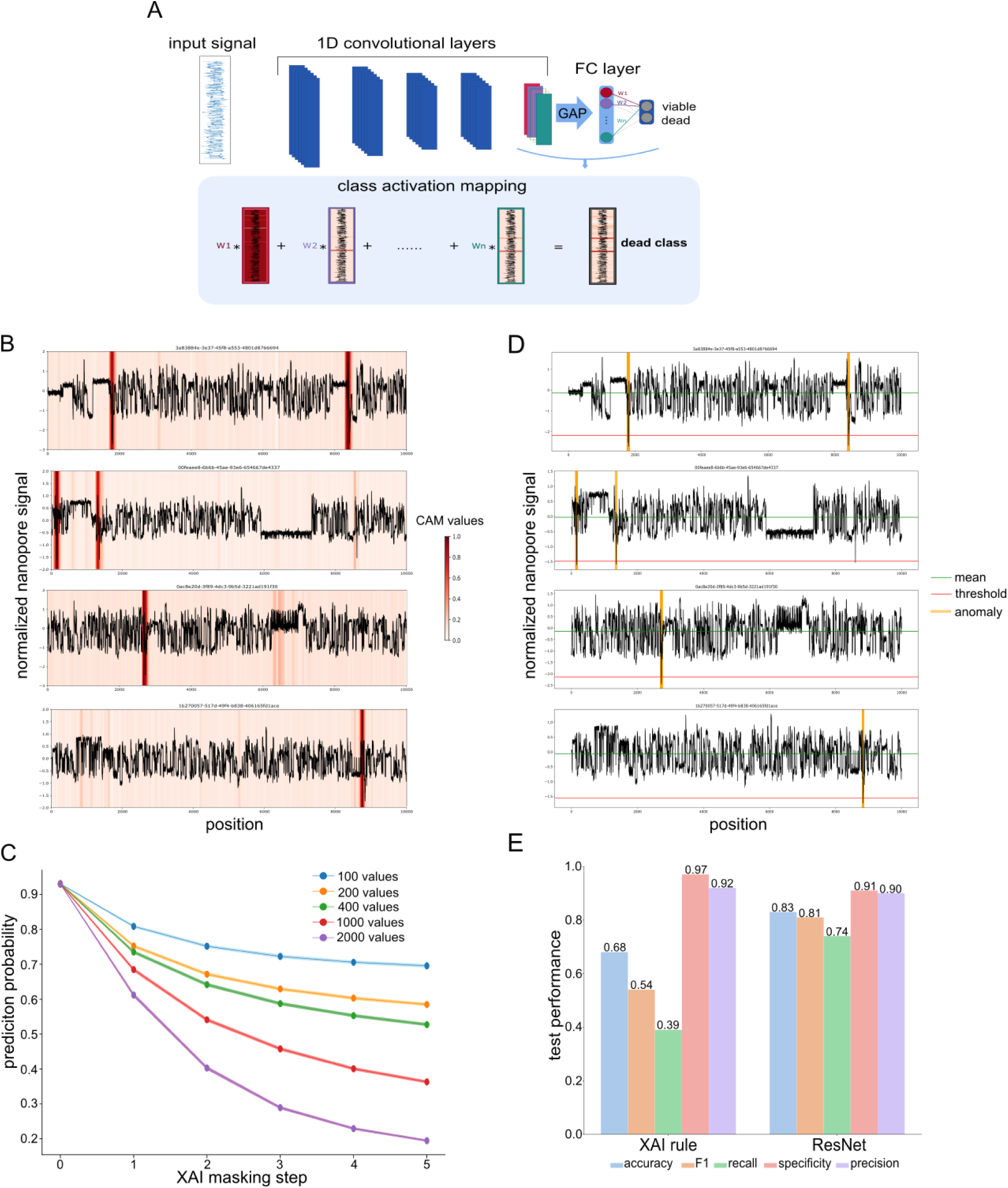
**Explainable AI for interpretability of ResNet1** (A) Class Activation Maps (CAMs) leverage the global average pooling (GAP) layer right before the fully connected (FC) layers of the residual neural network to map model interpretability onto the input features; they are generated by aggregating the final convolutional layer’s feature maps through a weighted sum, highlighting nanopore signal regions that allow the neural network to make accurate predictions. (B) Exemplary nanopore signal chunks that were classified as “dead” at a prediction probability of p>0.99, and their CAM values. Higher CAM values indicate stronger feature map activations. (C) Impact of consecutive masking (n=5 masking events) of the signal region with the highest CAM value per signal chunk (x-axis) on the model’s prediction probability (y-axis); five different mask sizes (from 100 to 2k signals) were used. (D) Application of a simplified XAI rule that classifies signal chunks according to the presence of a “sudden drop” (Materials and Methods; green: mean signal per chunk; red: threshold for sudden drop definition; yellow: identification of sudden drops in the exemplary signal chunks). (E) Comparison of the performance of the full model (ResNet1) with the simplified XAI rule.

We used the CAM regions to manually investigate the squiggle signals and found that many CAM regions of “dead” signal chunks included a sudden substantial drop in the nanopore signal. We therefore developed a simple algorithm that identifies such sudden drops, and applied this XAI rule to our test dataset (Materials and Methods; **Fig 2D**). We here classified any signal chunk with at least one sudden drop as “dead”, and all others as “viable”. While this simplified algorithm led to a drop in overall performance, we could still reach a relatively good overall accuracy of 0.68 (in comparison to 0.83 of the full model; **Fig 2E**). While the XAI rule maintained performance in terms of specificity and precision, we observed a substantial drop in recall in comparison to the full model (now 0.39 instead of 0.74). This shows that while the absence of a sudden drop in the nanopore signal data seems to reliably predict viability, not all “dead” signal chunks contain such a sudden drop. While this sudden-drop detection still seems to be at the core of our model’s interpretability (when focusing on high-confidence true positive chunks at p>0.99, the recall increased to 0.68), the model seems to additionally detect more subtle patterns in the nanopore signal data which allow it to increase recall while maintaining specificity and precision.

Based on our previous experience with squiggle data analysis [43, 44], we hypothesize that the substantial sudden drops in nanopore signal might be caused by a twist or kink in the DNA backbone, for example from 6-4 photoproduct pyrimidine dimers. The drop would then mark the event of a pore getting blocked due to such damage. Such a twist could also lead to a stalling signal if it impairs the motor protein from processing the DNA strand, which we indeed partially observed in our data (e.g., top signal chunk of **Fig 2B/D**). While we here hypothesize that UV exposure might have caused such twists in the DNA backbone, we intend to explore the biological, chemical, and physical features detected by squiggle-based viability models in more detail in future nanopore-based microbial studies.

### Sequencing read-level viability predictions

We finally assessed the performance of our viability model on the sequencing read-level instead of on the chunk-level by leveraging the capability of ResNets to handle variable input lengths. When shifting the test dataset analysis from the chunk- to the read-level (Materials and Methods), the prediction performances increased substantially, from an accuracy of 0.83 (chunk) to 0.96 (read), and with an improved AUPR of 0.99 (instead of 0.92 on the chunk-level) and AUROC of 0.99 (instead of 0.90 on the chunk-level) (**Fig 3A**; **Table S2**). These improvements indicate that the model might be able to use cumulative information per sequencing read to increase overall prediction performance. Additionally, such read-level viability predictions enable inferences from short sequencing reads that had to be excluded from chunk-level analyses; our model achieved good prediction performance on all such previously excluded short reads (in our case, reads shorter than <11.5k signals; n=166,628 reads; accuracy: 0.80, AUPR: 0.89, AUROC: 0.87). Given this improved performance, including on previously excluded short sequencing reads, and given that any genomic analysis including taxonomic assignment is usually applied to the unit of the read, we will report all prediction performances in the remaining manuscript on the level of sequencing reads.

**Fig 3.**
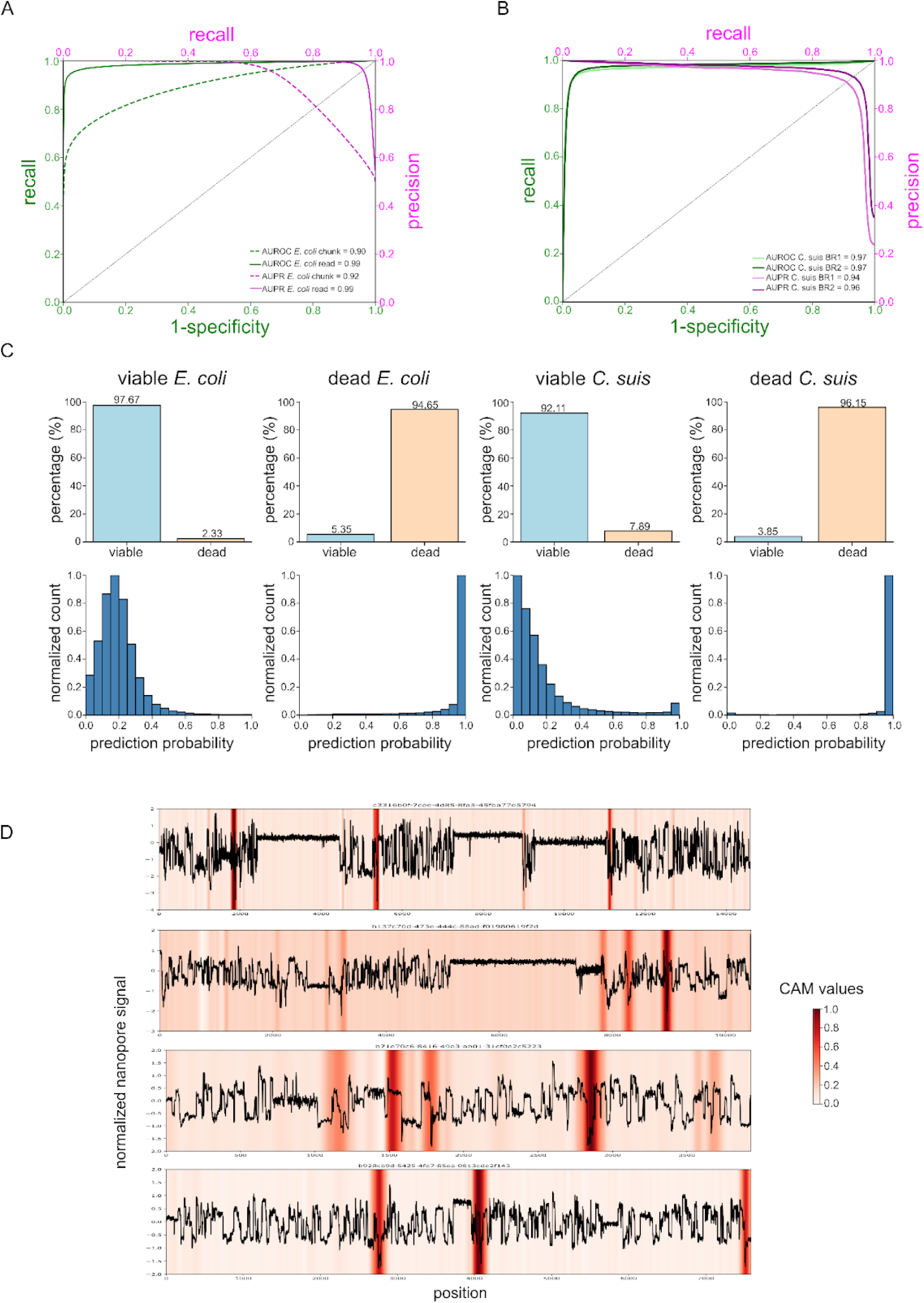
**Application of the *E. coli*-trained ResNet1 to obligate intracellular *Chlamydia suis*** (A) ResNet1 performance on the *E. coli* test dataset on the chunk-level (*dashed*) and on the sequencing read-level (*solid*) in terms of Precision-Recall (PR; *magenta)* and Receiver Operating Characteristic (ROC; *green*) curves and their respective Areas Under the Curve (AUPR, AUROC). (B) ResNet1 performance on the *C. suis* sequencing reads across two biological replicates (BR1 and BR2; *light* and *dark* lines, respectively) in terms of PR (*magenta)* and ROC (*green*) curves and their respective AUPR and AUROC. (C) Sequencing read-level comparison of ResNet1 classifications of the *E. coli* test and the *C. suis* datasets. *Top row*: Binary model predictions for viable and dead *E. coli* and *C. suis*, respectively, at the optimized prediction probability threshold of 0.5; *bottom row*: respective normalized distributions of model prediction probabilities across all sequencing reads. For *E. coli*, all 59,171 “viable” and 72,207 “dead” sequencing reads from the test dataset are shown; for *C. suis*, all 54,853 “viable” and 23,073 “dead” sequencing reads from both biological replicates are shown. (D) Squiggle signal of exemplary *C. suis* sequencing reads that were correctly classified as “dead”, and their CAM values; higher CAM values indicate stronger feature map activations (Materials and Methods).

### *Application to obligate intracellular* Chlamydia

We next applied our computational viability framework to distinguish viable from dead *Chlamydia suis* cells, an obligate intracellular bacterial species found endemically in the gastrointestinal tract of pigs with high infection rates in pig farms [45, 46]. Like other members of the *Chlamydiacea* family, these bacteria are defined by a complex biphasic life cycle comprising infectious elementary bodies and dividing reticulate bodies [47]. These unique properties render both cultivation- and vPCR-based approaches for viability estimations complicated [48]. We here took two samples from a viable *C. suis* culture as biological replicates (BRs 1 and 2), and subjected them to UV treatment (Materials and Methods). We obtained first insights into their viability using cultivation- and vPCR-based approaches (Materials and Methods). In the case of cultivation, UV-treated *C. suis* cells were unable to form viable inclusions in cell culture, whereas the untreated samples showed high infectivity with 8.48e7 (BR1) and 6.23e7 (BR2) inclusion forming units per mL (IFU/mL; **Table S3**). In the case of vPCR, all samples were processed with and without PMA [48] to quantify the respective amounts of chlamydial DNA with a sensitive *C. suis*-specific qPCR [49, 50] (Materials and Methods). We assessed the difference in copy number per mL between PMA-treated and untreated DNA as Δlog10, resulting in 0.82 and 0.81 for the viable samples, and 1.51 and 2.02 for the killed samples (BR1 and BR2, respectively; **Table S3**). These results are comparable to a previous study in which fresh *C. trachomatis* culture was heat-killed and an absence of viable *Chlamydia* resulted in a Δlog10 *Chlamydia* of 3.01 whereas a viability ratio of 100% resulted in a Δlog10 *Chlamydia* of 0.37 [48]. Cultivation and vPCR therefore confirmed that UV treatment had completely inactivated previously viable *C. suis* cells and had strongly reduced the amount of “viable” DNA in both biological replicates. Nanopore sequencing of the viable and dead *C. suis* cells identified 28,210 viable and 8,771 dead (BR1) and 26,643 viable and 14,302 dead (BR2) *Chlamydia*-classified sequencing reads (Materials and Methods; **Table S3**).

Our *E. coli* data-trained ResNet1 model achieved strong sequencing read-level viability prediction performances across both *C. suis* biological replicates (Materials and Methods). At the previously optimized probability threshold of 0.5, the model achieved an accuracy of 0.93 (both BRs), F1 score of 0.87 and 0.90 (BR1 and BR2), precision of 0.80 and 0.85, recall of 0.95 and 0.96, and specificity of 0.92 and 0.91, respectively (**Table S2**). Using threshold-independent metrics, the model achieved an AUROC of 0.97 (both BRs) and an AUPR of 0.94 and 0.96, respectively (**Fig 3B**). The slightly lower precision obtained by the model’s application to BR1 might stem from a slightly higher ratio of viable reads in this biological replicate (76.3% in BR2 *versus* 65.1% in BR2). When pooling the model’s viability predictions across the two biological replicates, the percentage of correctly classified sequencing reads (**Fig 3C**; *top row*) and the prediction probability distributions across reads (**Fig 3C**; *bottom row*) are comparable with sequencing read-level performances in the original *E. coli* test dataset (**Fig 3C**; *left*). This shows a certain degree of generalizability of our viability model beyond taxonomic boundaries despite the model being trained only on *E. coli* data. While both bacterial species are Gram-negative, they substantially differ in their ecology and life cycles, suggesting a potential applicability of deep models to nanopore squiggle data for taxonomy-agnostic viability predictions. The application of XAI to the *C. suis* sequencing reads further shows that the CAMs also highlight sudden drops in the *C. suis* squiggle data (**Fig 3D**), suggesting that UV exposure led to similar damage in both bacterial species. The killing method or, more generally, the source of degradation might therefore be the main determinant of viability-predictive features in nanopore squiggle data.

### Viability inference after antibiotic exposure of a mock community

To test whether our ResNet1 model that was trained on UV-killed *E. coli* could be used for viability predictions using a a different killing method, we generated two biological replicates (BR1 and BR2) of a mock community of carbenicillin-susceptible *E. coli* and carbenicillin-resistant *Klebsiella oxytoca*. Nanopore sequencing of the mock community after carbenicillin exposure resulted in 239,316 and 240,178 *E. coli* sequencing reads (number of reads mapping to the *Escherichia* genus in BR1 and BR2), and in 991,019 and 954,602 *K. oxytoca* sequencing reads for the experiment (number of reads mapping to the *Klebsiella* genus in BR1 and BR2; Materials and Methods). Across both species and biological replicates, the majority of the sequencing reads were classified as viable (*E. coli*: 82.2% and 76.4%; *K. oxytoca*: 80.8% and 87.4%, for BR1 and BR2, respectively), showing that ResNet1 could not distinguish susceptible from resistant bacteria after antibiotic exposure. This finding supports our hypothesis that the killing method or, more generally, the source of degradation might be the main determinant of viability-predictive features in nanopore squiggle data.

We therefore explored if we could train a new model using the same previously optimized ResNet1 architecture to accurately predict viability after antibiotic exposure. We first generated a clean antibiotic exposure training dataset of a single bacterial species by nanopore-sequencing the susceptible *E. coli* strain before and after exposure; similar to the UV-treated *E. coli*, we additionally made sure to (i) only sequence cell-free DNA for the dead viability class by stringent centrifugation and filtering of the supernatant, and to (ii) mainly sequence DNA from intact cells for the viable class by only processing the pellet after centrifugation (Materials and Methods). The newly trained antibiotic ResNet1 model achieved a test dataset accuracy of 0.73 on the sequencing read-level, with an AUPR of 0.98 and an AUROC of 0.87, and a performance on a heldout biological replicate of 0.68 accuracy, with an AUPR of 0.95 and an AUROC of 0.80 (based on 16,810 viable and 103,120 dead sequencing reads that were classified as *Escherichia*; **Table S4**; Materials and Methods).

We then applied this antibiotic ResNet1 model to the antibiotic exposure mock community; this time, the majority of sequencing reads from the susceptible *E. coli* were classified as dead (75.7% and 75.9%, for BR1 and BR2), while the majority of *K. oxytoca* was still correctly classified as viable (70.1% and 71.7%). The application of our XAI framework showed that this new antibiotic ResNet1 did not detect any sudden drops in the squiggle data that were previously identified by the UV ResNet1’s CAMs in dead sequencing reads (**Fig S4** for a few examples), showing that this new model seemed to have identified different squiggle signatures indicative of degradation through antibiotic exposure. These preliminary results suggest that our viability model can be tuned to classify viability in different degradation contexts, and that such models can potentially be applied to separate resistant from susceptible bacteria in mixed microbial communities.

### AI- and nanopore-empowered viability-resolved metagenomics

While metagenomic approaches provide the unique opportunity of generating *de novo* assemblies and potentially complete microbial genomes to explore the “microbial dark matter” as well as to infer potential functions such as metabolic and virulence potential [8, 9], they have suffered from their inability to differentiate between viable and dead microorganisms [2, 15]. Such viability inferences can, however, distort any microbial inference, ranging from assessing ecosystem functions of environmental microbiomes to inferring the virulence of potential pathogens. As established viability-resolved metagenomic approaches are labor-intensive as well as biased and lack sensitivity (e.g., <u>23</u>), we here show first evidence that a fully computational framework based on residual neural networks with convolutional data processing layers can leverage raw nanopore signal data, also known as squiggle data, to make accurate inferences about microbial viability. Using experimentally killed bacterial cultures and a simple mock microbial community, we show that such models can infer viability from sequencing reads at high accuracy, potentially allowing for simultaneous taxonomic and viability classifications in metagenomic datasets.

We first leverage microbial degradation through UV exposure to show that such viability models can make accurate predictions in a taxonomy-agnostic manner; a model that has only ever seen *E. coli* squiggle data can predict viability in UV-exposed *Chlamydia* at high recall and specificity (>0.9). The application to estimate the viability of pathogenic *Chlamydia* is hereby of potentially immediate interest to veterinary scientists since traditional methods for assessing the pathogen’s viability have been labor-intensive and suffered from inherently high false negative rates; more research is, however, needed to assess to what extent this model can capture natural degradation in *Chlamydia*.

Our subsequent XAI analyses point to the potential role of UV-induced DNA backbone damage for achieving accurate model predictions in both species; the simplified XAI rule is, however, not sufficient to correctly classify the majority of “dead” signal chunks (recall of 0.39), which means that the residual neural network has apparently identified additional more subtle signal patterns that allow the full model to make more sensitive predictions (recall of 0.74) while maintaining specificity and precision. More research is therefore needed to fully understand the biological, physical, or chemical underpinnings of our viability model predictions; we, however, anticipate that future experiments for squiggle data generation, including on different taxonomic groups and killing methods, will help us tease apart the origins of our current XAI results.

Besides exploring the underlying rules of squiggle signal patterns, the generalizability of such squiggle-based viability models also needs to be assessed in more detail, including for a wider breadth of microbial taxa such as spore-forming bacteria or fungi, and for a variety of degradation sources and intensities. Our preliminary results indicate that our deep model that can accurately predict UV-induced degradation can not distinguish susceptible and resistant bacteria in an antibiotic exposure experiment of a simple mock community. We, however, show that training the same residual neural network architecture on newly generated killing method-specific squiggle data can achieve good accuracy in the respective mock community (>0.7). While this application shows the limits of our current models’ generalizability, we also argue that clinical metagenomics might already profit from such antibiotic-specific viability inferences: As previously discussed, certain disinfection methods or systemic antibiotics in the clinical setting often kill the bacteria before the DNA is destroyed [15, 17], leading to potential false-positive pathogen detections using metagenomics; our viability model could give additional information on the antibiotic exposure’s impact, and confidence could be increased by accumulating viability evidence across sequencing reads per pathogen of interest. In addition, we hypothesize that models could be trained to detect damage induced by suboptimal antibiotic usage, which has been implicated in the emergence of new antibiotic resistances [51].

In order to achieve true metagenomic applications in the future, the training data of a single model could be diversified in terms of taxonomy and degradation, potentially increasing the generalizability of its viability inferences. We also anticipate that quantitative AI modeling has the potential to inform more differentiated viability assessments, which might help quantify or time degradation events, and even decipher the relevance of dormancy and metabolic inactivity in metagenomic studies [52, 53]. We envision many potential benefits of such a widely applicable computational framework for microbial viability inference, including for applications in environmental, veterinary, and clinical settings. As is the case for epigenetic inferences [36, 37, 38], the viability inference-enabling squiggle data is a complementary output of any nanopore sequencing experiment of native DNA (and RNA), that is then usually basecalled and archived for future re-basecalling after potential basecalling model improvements. This means that any future nanopore-based metagenomic study could make viability predictions for free without additional costs and laboratory work, and that any existing archived nanopore data could be assessed in terms of its microorganisms’ viability—which would allow us to quantify the impact of dead microorganisms on metagenomic analyses in a diversity of datasets and ecological settings, and to further explore factors such as species- and environment-specificity.

## Materials and Methods

### Viability inference explorations in UV-exposed E. coli

1. *Training data generation*

We cultured *E. coli K12* in 200 mL Luria-Bertani (LB) medium for 24 hours at 37°C to reach the log phase of the growth curve. The culture was then used to inoculate four 200 mL LB media in 1 L Erlenmeyer flasks, which were again incubated for 24 hours to reach the growth log phase. One of the media was used as viable control, i.e. DNA was extracted from 750 uL of the living culture using the spin-column-based QIAGEN PowerSoil Pro Kit (QIAGEN, 2018, Hilden, Germany), following the manufacturers’ instructions. The remaining three cultures were killed by one of the following stressors: UV irradiation at 254 nm for 15 min, heat shock at 120°C for 5 min, or bead beating for 30 min. To then separate extracellular DNA from cell debris and intact bacterial cells, we centrifuged the media for 10 min at 4,000 x g and filtered the supernatant through 0.2 µm filters. The resulting extracellular DNA was subsequently kept at room temperature for 5 days to simulate the natural accumulation of DNA degradation. DNA from dead bacteria was extracted from these samples using the same extraction approach following the QIAGEN PowerSoil Pro Kit protocol, but the first lysis buffer step was omitted since cell lysis had already happened.

We then used Oxford Nanopore Technologies’ Rapid Barcoding library preparation kit (RBK114-24 V14), R10.4.1 MinION flow cells, and MinKNOW software v23.04.5 for shotgun nanopore sequencing of the “viable” and “dead” DNA. We used four barcodes for each sample, resulting in DNA input of 800 ng and 218 ng for the preparation of the “viable” and “dead” library, respectively. We ran each library for 24 h, using two different flow cells to avoid any cross-contamination, and filtered the resulting nanopore data at a minimum read length of 20 b. Raw nanopore data was created using the standard translocation speed of 400 b/s and a sampling frequency of 5 kHz. We applied Dorado (https://github.com/nanoporetech/dorado) SUP-basecalling model v4.2.0 (dna_r10.4.1_e8.2_400bps_sup@v4.2.0) and 6mA-aware SUP-basecalling (6mA@v1) to all nanopore reads that had passed internal data quality thresholds to obtain *E. coli* DNA sequence data. We subsequently removed sequencing adapters and barcodes using Porechop v0.2.3 (https://github.com/rrwick/Porechop).

1. *Model training*

We tested the implementation of different residual neural networks and transformer architectures to predict the binary viability state from the raw nanopore data (0=viable; 1=dead). The first residual neural network, ResNet1, consists of four layers, each containing two bottleneck blocks. Each bottleneck block consists of convolutional layers, batch normalization, and a rectified linear unit (ReLu) activation function. Each of the four layers consists of an increasing number of convolutional channels: 20, 30, 45, and 65, respectively, followed by global average pooling and a fully connected layer, resulting in 66,916 parameters. The model then uses a softmax function to convert logits, the raw outputs from the fully connected layer, into predicted probabilities ranging from 0 to 1. We evaluated the training of the model using the Adam optimizer for mini-batch gradient descent at three different LRs (1e-3, 1e-4, and 1e-5), training the model up to 1,000 epochs and at a batch size of 1,000 signal chunks. We initialized the model using Kaiming initialization. For ResNet2, we increased the number of convolutional channels to 40, 60, 90, and 135, respectively, resulting in 1,828,777 parameters. For ResNet3, we increased the number of convolutional channels to 512, 30, 45, and 67, respectively, resulting in 2,479,140 parameters. The transformer model was based on a positional encoding, a convolutional layer with a channel number of 24, and one block of one attention head, resulting in 219,586 parameters.

We processed the *E. coli* squiggle data by excluding the first 1,500 signal points (potential noise, adapter sequences, or barcodes), then cutting it into signal chunks of 10k signals, and separated the chunks into balanced training (60%), validation (20%), and test (20%) set along each original sequencing read to avoid that signal chunks from the same read would end up in the same dataset. We pooled the viable and dead signal chunks to obtain exactly balanced training, validation, and test sets. For normalizing each chunk, we subtracted the median per chunk and divided it by the median absolute deviation (MAD) to make the signal data robust to outliers. We then scaled the signal by the MAD scaling factor 1.4826 and replaced outliers exceeding 3.5 times the scaled MAD by the mean of their two neighboring values.

We also trained ResNet1 on the basecalled nanopore data (with or without 6mA basecalling) of viable and dead *E. coli* at a standardized chunk size of 800 b, which roughly corresponds to a signal chunk size of 10k signals. For encoding, we used a one-hot encoding method to turn DNA sequences into unique binary vectors. We then concatenated and saved these encoded sequences as tensors for training and testing. We finally trained ResNet1 on signal chunks of different signal lengths, ranging from 1k to 20k signals.

For logistic regression training, we used the LogisticRegression class from scikit-learn v1.2.2 with default parameters.

1. *Explainable AI*

We utilized CAMs to identify and visualize signals regions that influenced the model’s decision-making. As feature maps from the final convolutional layer undergo a global average pooling layer where each map is averaged and concatenated, we can calculate the weighted sum of these feature maps using the weights of the fully connected layer and project it back onto the preprocessed signal [42]. To do so, we implemented CAM in Python/PyTorch. During the forward pass, we ensure that the feature maps from the last convolutional layer are captured. To compute the CAM, we use the weights of the model’s output layer for the class of interest, multiplying these weights with the corresponding feature maps and then summing them up. We convert the resulting CAM to an array and normalize its values to a range of 0 to 1. We then overlay the CAM on the original input signal to identify the regions most influential in the model’s decision-making process. We additionally used the Remora API to match raw nanopore data to the corresponding Dorado-basecalled bases, to then manually investigate any obvious sequence abnormalities in the CAM regions.

For consecutive masking of the CAM regions with the highest CAM values, we used Python to first obtain and normalize the CAM values of all true-positive signal chunks at p>0.5 (n=286,179), identify the maximum value, mask the signal region (i.e., setting to zero after MAD-normalization) at a specified mask size (between 100 and 2k signals) centered around the maximum-CAM signal, and obtain updated prediction probabilities. We repeated this masking step five times and calculated the confidence interval at each masking step: We obtained the mean and Standard Error of the Mean (SEM) of the newly calculated prediction probabilities across all signal chunks to calculate the 95% confidence interval at mean ± 1.96 * SEM. We plotted the results using matplotlib.

We next used Python to develop an algorithm to obtain a simplified XAI rule to distinguish “dead” from “viable” signals chunks based on our CAM results by identifying the presence of at least one sudden drop in the chunk. To identify those sudden drops, we first calculated the mean and standard deviation (SD) of each signal chunk, and found that a scaling factor of 3 identified most manually detected sudden drops at a vertical threshold of mean - 3 * SD.

1. *Sequencing read-level inferences*

To evaluate read-level performance, we leveraged the inherent flexibility of ResNet architectures to handle variable-length input. Prior to all read-level analyses, we removed chimeric reads using information from Dorado basecalling. Dorado splits chimeric reads and tags the resulting child reads with a “pi” tag (parent_read_id) in the BAM file, indicating their origin from an unsplit read. Using the pysam Python package, we extracted the read IDs of the reads carrying the “pi” tag to exclude chimeric reads from our downstream inference analysis. The non-chimeric reads were subsequently filtered according to Kraken2 v2.1.4-based taxonomic assignment to the *Escherichia* genus [54]. All sequencing read-level inferences were made only on non-chimeric, correctly taxonomically classified (on the genus level) reads.

### *Application to obligate intracellular* Chlamydia

The *Chlamydia suis* strain S45 (kindly provided by J. Storz, Baton Rouge, LA, USA) was cultured as described by Leonard *et al.* [55]. Briefly, chlamydiae were grown in the epithelial rhesus monkey kidney cell line LLM-MK2 (provided by IZSLER, Brescia, Italy) and prepared as semi-purified stock by scraping infected cells into the supernatant and removal of cellular debris by centrifugation at 500 g for 10 min. Chlamydiae were then pelleted (10’000 x g, 45 min) and resuspended in sucrose phosphate glutamate buffer (SPG). To determine the viability of this stock, one aliquot was thawed on ice and separated into two tubes of which one was UV-inactivated using a Hoefer UVC 500 Ultraviolet Crosslinker (Hoefer Inc., Bridgewater MA 02324, United States): Briefly, samples were kept on ice and exposed to 8 watts of UV light at 12.5 cm for 30 min, similar to previously described protocols [56]. All samples were prepared in biological duplicates. UV-inactivated chlamydia were then further incubated at room temperature for 48 h prior to further processing, while viable chlamydiae were immediately processed. The resulting four samples were divided into subsamples and used for cultivation, vPCR, and nanopore sequencing.

For viability determination by culture, subsamples (approximately 1e7 IFU) were used to infect four glass coverslips (13 mm in diameter, ThermoScientific, Waltham, MA, USA) in 24-well plates (TPP Techno Plastic Product AG, Trasadingen, Switzerland) seeded to confluence with LLC-MK2 cells [57]. For the “viable” subsamples, a 1:1000 dilution was performed prior to inoculation. The infection was then enhanced by centrifugation for 1 h at 25°C (1000 x g). After 48 h of incubation at 37°C (5% CO₂), cultures were fixed for 10 min in ice-cold methanol. If cultures were free of chlamydiae, one well was scraped and transferred to fresh monolayers up to three times to confirm negativity as described [58]. Coverslips were then processed using a well-established immunofluorescence assay [59]. Briefly, DNA was stained with 1 μg/mL 4’, 6-diamidino-2’-phenylindole dihydrochloride (DAPI, Molecular Probes, Eugene, OR, USA). In parallel, inclusions were labeled with a *Chlamydiaceae*-specific primary antibody (*Chlamydiaceae* LPS, Clone ACI-P; Progen, Germany), which was diluted 1:200 in blocking solution consisting of 1% bovine serum albumin (BSA, St. Louis, MO, USA) in phosphate-buffered saline (PBS, GIBCO, Invitrogen, Carlsbad, CA, USA). Inclusions were visualized with Alexa Fluor 488 goat anti-mouse (Molecular Probes) diluted 1:500 in blocking solution. As a final step, coverslips were washed with PBS, mounted with FluoreGuard (Hard Set; ScyTek Laboratories Inc., Logan, UT, USA) on glass slides, and inclusions determined using a Leica DMLB fluorescence microscope (Leica Microsystems, Wetzlar, Germany) and a 10X ocular objective (Leica L-Plan 10x/ 25 M, Leica Microsystems). In parallel, a three-fold dilution series of the sample was performed in 96-well plates (TPP) and processed as above. The number of IFU/ml for the whole samples was then calculated based on the number of inclusions detected using the Nikon Ti Eclipse epifluorescence microscope (Nikon, Tokyo, Japan) at a 20X magnification [57].

For vPCR, subsamples (approximately 2e7 IFU) were taken and mixed with 200 µl SPG and 100 µl PMA enhancer for Gram Negative Bacteria (5X, Biotium, Fremont, CA, USA). PMAxx (Biotium) at a final concentration of 50 µM was added (“PMA-treated”) or not (“untreated”) to the subsamples. Samples were then exposed to a 650-W light source using a PMA-Lite LED Photolysis Device (Biotium) for 5 min, followed by 2 min on ice and additional light exposure for 5 min [48]. For vPCR as well as nanopore sequencing (subsamples with approximately 5e8 IFU), DNA was extracted using the DNeasy® Blood and Tissue Kit (QIAGEN, Hilden, Germany) according to the manufacturer’s instructions. The amount of chlamydial DNA was quantified with a sensitive *C. suis* qPCR [60], based on a standard curve with recombinant plasmid containing the amplicon target and calculated for the whole sample. For subsequent nanopore sequencing of the DNA extracts (viable: 13–16 ng/μL; dead: 1.9–2.8 ng/μL), we followed the same approach as described for *E. coli* above. Each viable sample was sequenced using one barcode (input volume of 10 μL) and each dead sample using three barcodes (input volume of 30 μL). Sequencing reads were processed following the same steps used for the *E. coli* dataset. Chimeric reads were identified using the ’pi’ tag added by Dorado and removed with the pysam Python package. Adapter and barcode trimming was performed using Porechop, and taxonomic classification was conducted with Kraken2 v2.1.4, retaining only reads classified as *Chlamydia*.

### Antibiotic exposure experiment

To generate data from a simple mock community, a carbenicillin-susceptible *E. coli* strain and an ESBL-producing *Klebsiella oxytoca* strain (ATCC 700324) were cultured on Müller-Hinton (MH) agar at 37 °C for 20 hours in two biological replicates. Susceptibility to penicillin as an approximation of carbenicillin resistance was confirmed by VITEK2 and growth curve analysis at 100 and 200 ng/μL. Species identity was confirmed by MALDI-TOF MS and MLST. Cultures were inoculated at OD600 = 0.01 in 10 mL MH broth and incubated for 5 hours. Carbenicillin (200 ng/μL) was added during logarithmic growth, followed by 20 hours of further incubation. DNA was extracted directly from 250 μL of uncentrifuged culture using the same DNA extraction approach as described for UV-exposed *E. coli*.

To generate clean training data for an antibiotic exposure-specific model, two biological replicates of the same *E. coli* strain were used. For the viable samples, DNA was extracted from centrifuged pellets (4000 × g, 10 min) before antibiotic treatment, respectively. For the dead samples, DNA was extracted from the filtered supernatant (0.2 μm) 20 hours after carbenicillin exposure using the same DNA extraction approach as described for UV-exposed *E. coli*.

Nanopore sequencing and data processing were done as described for *E. coli* above. All inferences were done on the sequencing read-level and restricted to non-chimeric reads classified as *Escherichia* or *Klebsiella*, respectively. Model training was also done as described above, using the optimized ResNet1 architecture, but chunk size was set to 5k instead of 10k signals to increase the size of the training dataset. The final dataset consisted of 330,000 chunks for training, 110,000 for validation, and 110,000 for testing. The model was trained for up to 600 epochs using the Adam optimizer with a learning rate of 1e-5. The best validation accuracy was achieved at epoch 550.

## Data Availability Statement

All raw data has been made publicly available via ENA (study accession number: PRJEB76420). All code and the UV ResNet1 model (“UV_ecoli_ResNet_677ep.ckpt”) and antibiotic ResNet1 model (“antibiotic_ecoli_ResNet_550ep.ckpt”) have been made publicly available via Github: https://github.com/Genomics4OneHealth/Squiggle4Viability.git.

## Acknowledgments

We thank the laboratory of the Research Unit of Comparative Microbiome Analysis at Helmholtz Munich, Germany, especially Cornelia Galonska, for their support in processing the *Escherichia coli* samples. We thank the laboratory of the Institute of Veterinary Pathology, Switzerland, especially Theresa Pesch, for their support in processing the *Chlamydia suis* samples. We further thank Valentin Rauscher for his help in training several deep models during his internship in the Urban research group at Helmholtz Munich.

## Supporting information

**Table S1.** UV-killed *E. coli* viability inferences of deep neural network and logistic regression models. Test dataset performance metrics of residual neural networks (“ResNet”), transformer architectures, and logistic regression models, trained on various data modalities (nanopore squiggle “Signal” or basecalled DNA “Nucleotide” sequence) at various signal chunk sizes (“Length”), and on sequencing read length or translocation speed (“Trans speed”).

| Model<br>Architecture | Data<br>Modality | Length | Accuracy | F1 | Precision | Sensitivity | Specificity | AUROC | AUPR |
| --- | --- | --- | --- | --- | --- | --- | --- | --- | --- |
| ResNet1 | Signal | 10K | 0.83 | 0.84 | 0.78 | 0.91 | 0.74 | 0.90 | 0.87 |
| ResNet2 | Signal | 10K | 0.83 | 0.85 | 0.77 | 0.94 | 0.72 | 0.89 | 0.86 |
| ResNet3 | Signal | 10K | 0.81 | 0.82 | 0.76 | 0.89 | 0.72 | 0.87 | 0.83 |
| Transformer | Signal | 10K | 0.79 | 0.82 | 0.73 | 0.92 | 0.67 | 0.86 | 0.82 |
| ResNet1 | Nucleotide<br>(A,C,G,T) | 800 | 0.51 | 0.51 | 0.51 | 0.52 | 0.50 | 0.52 | 0.53 |
| ResNet1 | Nucleotide<br>(A,C,G,T, M) | 800 | 0.51 | 0.50 | 0.51 | 0.50 | 0.52 | 0.51 | 0.51 |
| ResNet1 | Signal | 1K | 0.58 | 0.65 | 0.55 | 0.80 | 0.35 | 0.61 | 0.58 |
| ResNet1 | Signal | 5K | 0.71 | 0.76 | 0.66 | 0.89 | 0.54 | 0.78 | 0.73 |
| ResNet1 | Signal | 7K | 0.77 | 0.79 | 0.72 | 0.88 | 0.65 | 0.83 | 0.79 |
| ResNet1 | Signal | 12K | 0.84 | 0.85 | 0.77 | 0.97 | 0.71 | 0.91 | 0.88 |
| ResNet1 | Signal | 20K | 0.89 | 0.90 | 0.85 | 0.95 | 0.83 | 0.94 | 0.91 |
| Regression | Read length | NA | 0.50 | 0.00 | 0.00 | 0.00 | 1.00 | 0.56 | 0.56 |
| Regression | Trans speed | NA | 0.59 | 0.55 | 0.61 | 0.50 | 0.68 | 0.63 | 0.59 |

**Table S2.** Sequencing read-level viability inferences of UV ResNet1. Performance metrics across sequencing reads of the ResNet1 model trained on UV-killed *E. coli* for the *E. coli* test dataset, and two biological replicates (BR1 and BR2) of UV-killed and viable *Chlamydia suis*. The number of total reads is the number of sequencing reads after processing of the nanopore sequencing data by Porechop; the number of genus-classified reads is the number of sequencing reads that map to the *Escherichia* or *Chlamydia* genus, respectively, using Kraken2 (Materials and Methods).

| Metric | *E. coli* test dataset | *C. suis* BR1 | *C. suis* BR2 |
| --- | --- | --- | --- |
| #total reads | 140,268 | 63,935 | 76,418 |
| #genus-classified reads | 131,378 | 36,981 | 40,945 |
| Accuracy | 0.96 | 0.93 | 0.93 |
| F1 Score | 0.96 | 0.87 | 0.90 |
| Sensitivity | 0.94 | 0.80 | 0.85 |
| Specificity | 0.97 | 0.95 | 0.96 |
| Precision | 0.98 | 0.92 | 0.91 |
| AUPR | 0.99 | 0.97 | 0.97 |
| AUROC | 0.98 | 0.94 | 0.96 |

**Table S3.** Cultivation, viability PCR (vPCR) and nanopore sequencing metrics of UV-killed *Chlamydia suis*. Cultivation titer in number of Inclusion Forming Units (IFUs) per mL; PMA-untreated vPCR reflecting total *Chlamydia* content, PMA-treated vPCR reflecting viable *Chlamydia* content, and Δlog10 of the PMA-treated copy number divided by the PMA-untreated copy-number reflecting overall viability; total number of nanopore sequencing reads after Porechop-processing, and number of *Chlamydia*-classified reads using Kraken2 (Materials and Methods), of two biological replicates (BR1 and BR2) of viable and dead *C. suis*.

| Condition | Titer [IFU/mL] | PMA-untreated vPCR [copy number per mL] | PMA-treated vPCR [copy number per mL] | $\Delta\log_{10}$ of viability ratio | #total reads | #genus-classified reads |
| --- | --- | --- | --- | --- | --- | --- |
| viable BR1 | 8.48e<sup>7</sup> | 9.63e<sup>7</sup> | 1.47e<sup>7</sup> | 0.82 | 37,439 | 28,210 |
| dead BR1 | 0 | 7.16e<sup>5</sup> | 2.23e<sup>4</sup> | 1.51 | 26,496 | 8,771 |
| viable BR2 | 6.23e<sup>7</sup> | 9.15e<sup>7</sup> | 1.24e<sup>7</sup> | 0.81 | 34,444 | 26,643 |
| dead BR2 | 0 | 1.03e<sup>6</sup> | 9.81e<sup>3</sup> | 2.02 | 41,974 | 14,302 |

**Table S4.** Sequencing read-level viability inferences of antibiotic exposure ResNet1. Performance metrics across sequencing reads of the ResNet1 model trained on antibiotic-exposed *E. coli* for the *E. coli* test dataset, and a heldout biological replicates (BR). The number of *E. coli* reads is the number of sequencing reads after processing of the nanopore sequencing data by Porechop and mapping to the *Escherichia* genus using Kraken2 (Materials and Methods).

| Metric | Test dataset | BR |
| --- | --- | --- |
| #*E. coli* reads | 49,781 | 119,930 |
| Accuracy | 0.73 | 0.68 |
| F1 | 0.82 | 0.78 |
| Sensitivity | 0.71 | 0.66 |
| Specificity | 0.87 | 0.80 |
| Precision | 0.98 | 0.95 |
| AUPR | 0.98 | 0.95 |
| AUROC | 0.87 | 0.80 |

**Fig S1.**
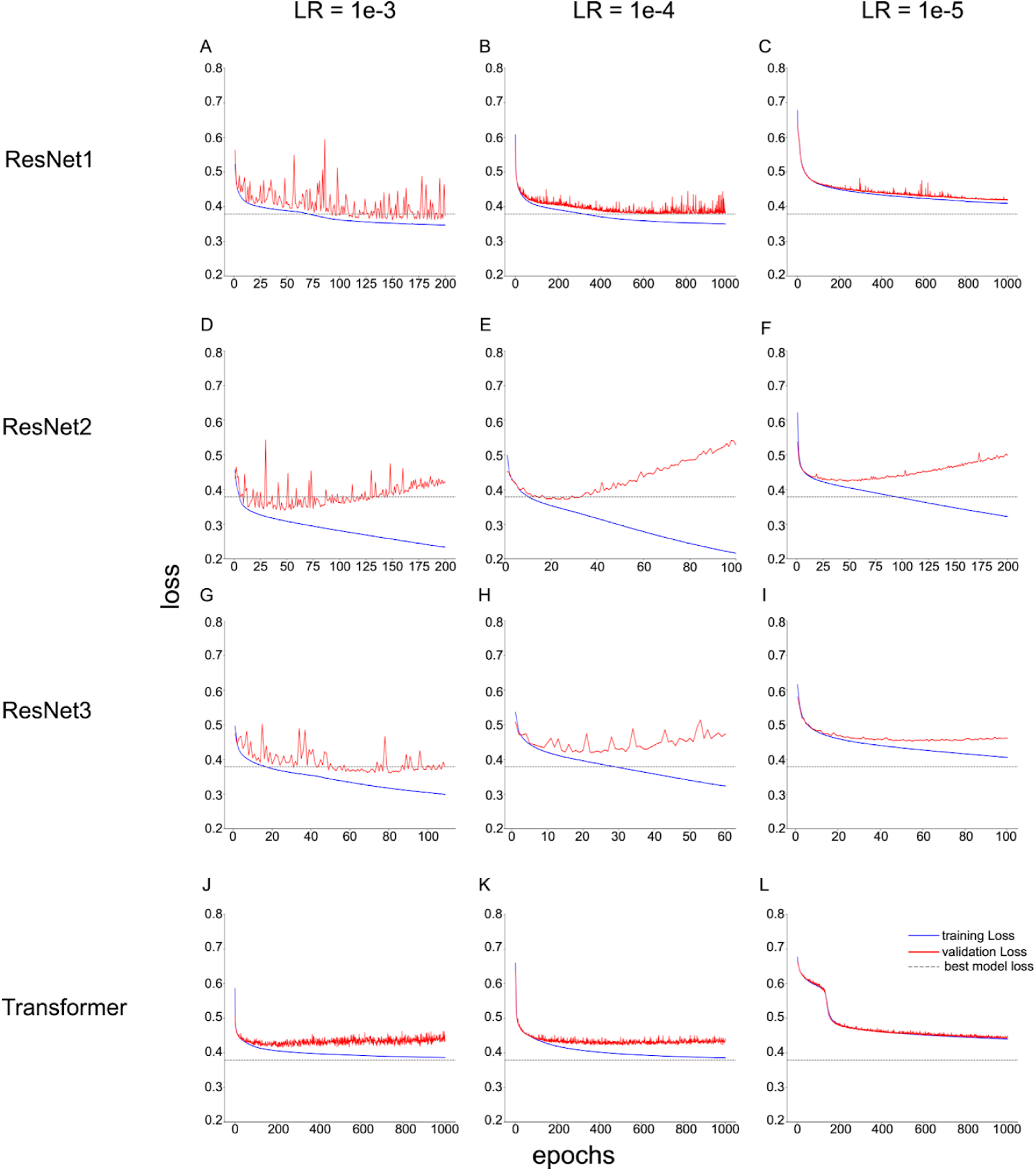
Training and validation loss across deep neural network architectures tested for nanopore squiggle signal-based viability inference. (A-C) Model loss of ResNet1 at learning rates (LRs) of 1e-3, 1e-4, and 1e-5; (D-F) model loss of ResNet2 at LRs of 1e-3, 1e-4, and 1e-5; (G-I) model loss of ResNet3 at LRs of 1e-3, 1e-4, and 1e-5; and (J-L) model loss of the transformer models at LRs of 1e-3, 1e-4, and 1e-5. The solid blue line indicates the training loss, the solid red line indicates the validation loss, and the dashed line indicates the minimum validation loss from the final ResNet1, LR=1e-4, model.

**Fig S2.**
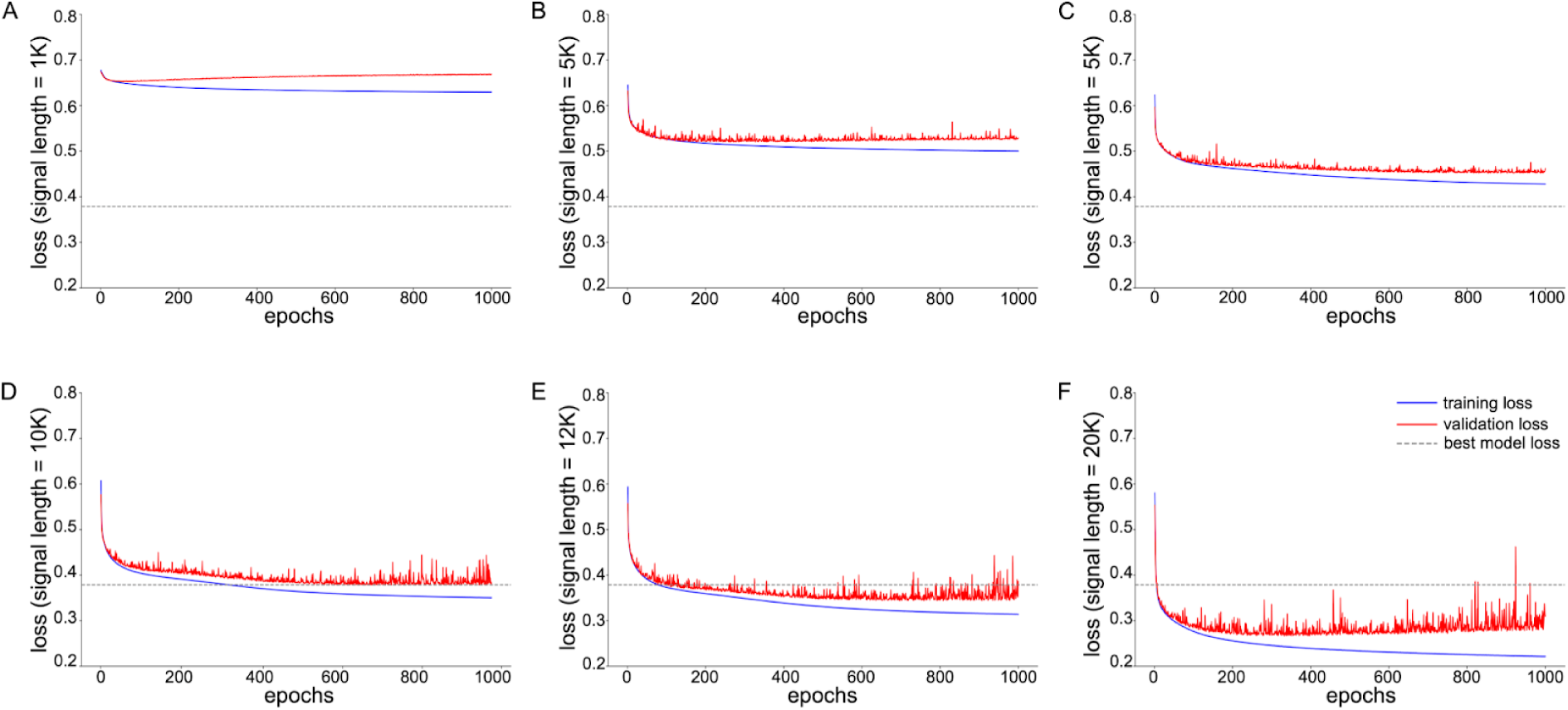
Training and validation loss of ResNet1 at various signal chunk sizes. The signal chunk size varies from (A) 1k, (B) 5k, (C) 7k, (D) 10k, to (E) 12k and (F) 20k. The solid blue line indicates the training loss, the solid red line indicates the validation loss, and the dashed line indicates the minimum validation loss from the final ResNet1model using a signal chunk size of 10k.

**Fig S3.**
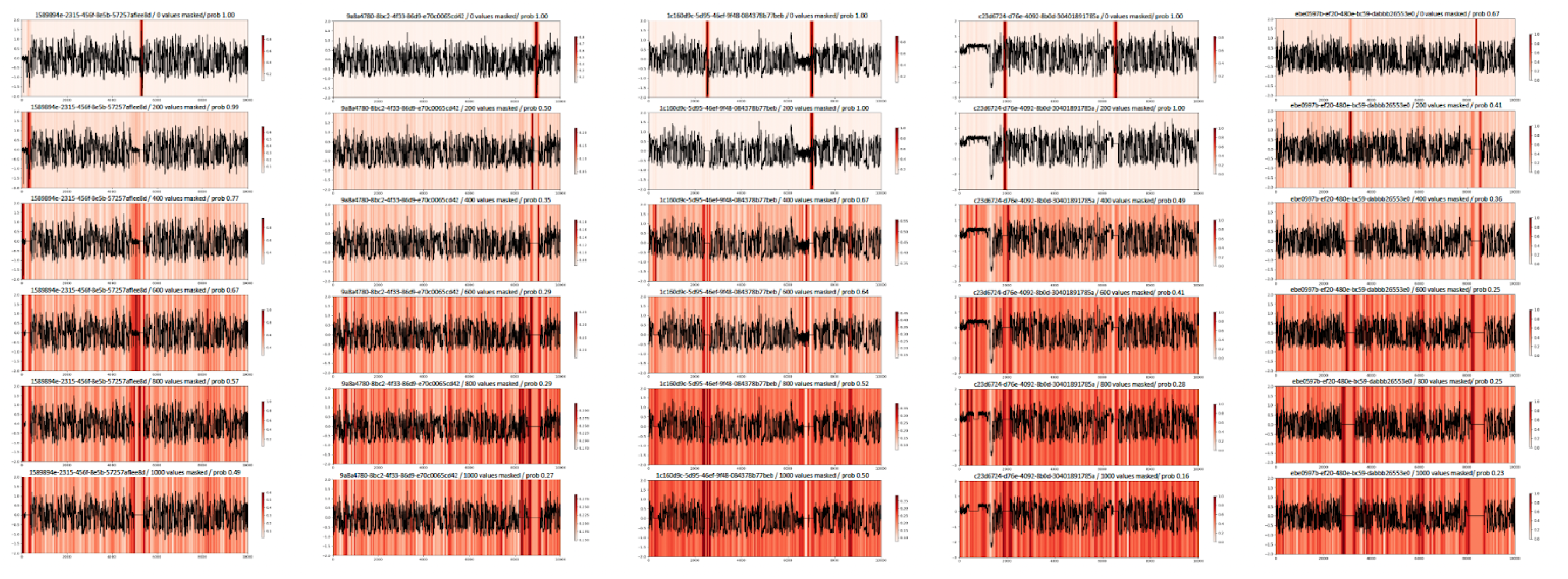
Exemplary drops in ResNet1 prediction probabilities in nanopore signal chunks after consecutive masking of the signal region with the respectively highest CAM value. *Figure headers*: signal chunk ID / total number of masked signal values / prediction probability per signal chunk “prob”. *Left to right*: Five exemplary nanopore signal chunks (length of 10k signals). *Top to bottom*: Consecutive masking of 200 signal values per masking event (Materials and Methods). *Legends*: Red-colored CAM value visualizations; higher CAM values indicate stronger feature map activations.

**Fig S4.**
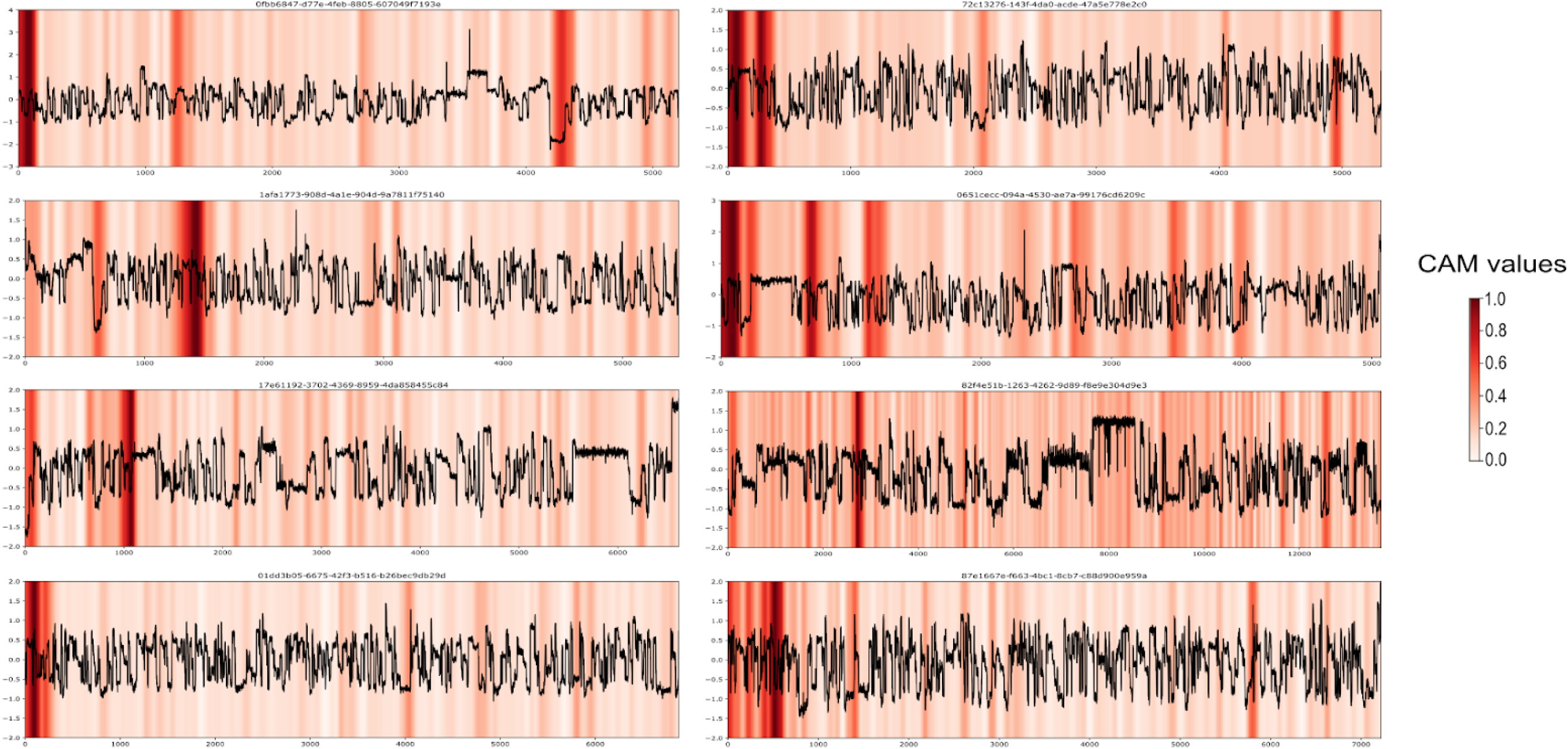
Exemplary nanopore signal patterns of antibiotic-killed *E. coli* sequencing reads and XAI Class Activation Mapping (CAM). *Legend*: Red-colored CAM value visualizations; higher CAM values indicate stronger feature map activations.

